# GPR41/43 regulates blood pressure by improving gut epithelial barrier integrity to prevent TLR4 activation and renal inflammation

**DOI:** 10.1101/2023.03.20.533376

**Authors:** Rikeish R. Muralitharan, Tenghao Zheng, Evany Dinakis, Liang Xie, Anastasia Barbaro-Wahl, Hamdi A. Jama, Michael Nakai, Madeleine Patterson, Chad Johnson, Ekaterina Salimova, Natalie Bitto, Maria-Kaparakis Liaskos, David M. Kaye, Joanne A. O’Donnell, Charles R. Mackay, Francine Z. Marques

**Author notes:** Corresponding author: A/Prof Francine Marques, Hypertension Research Laboratory, School of Biological Sciences, Faculty of Science, Monash University, Melbourne, Australia.

## Abstract

Fermentation of dietary fibre by the gut microbiota leads to the production of metabolites called short-chain fatty acids (SCFAs), which have emerged as potent regulators of immune, metabolic, and tissue barrier functions. More recently, a high fibre diet and SCFA supplementation were shown to lower blood pressure and be cardio-protective. SCFAs activate host signalling responses via the receptors GPR41 and GPR43, which have redundancy in their signalling pathways. Whether these receptors play a role in hypertension or mediate the cardio-protective effects of fibre remains unknown. Using an experimental model that lacks both GPR41 and GPR43, we show that lack of signalling via these receptors increases risk to high blood pressure and leads to cardiorenal fibrosis and hypertrophy.

Moreover, we demonstrate that GPR41/43 signalling is essential in maintaining gut epithelial barrier, which prevents the translocation of the bacterial toxins lipopolysaccharides (LPS) from entering the peripheral circulation. In the absence of GPR41/43, this is accompanied by macrophage infiltration to the kidneys, resulting in pro-inflammatory cytokine production.

Using an antagonist against the LPS’ receptor, TLR4, a potent pro-inflammatory signalling pathway, we were able to rescue the cardiovascular phenotype in GPR41/43 knockout mice. We also demonstrate that GPR41/43 are, at least partially, responsible for the blood pressure-lowering and cardio-protective effects of a high fibre diet; however, improvements of gut barrier integrity and macrophages in the kidney were independent of GPR41/43 signalling.

Finally, using the UK Biobank, we provide translational evidence that variants associated with lower expression of both GPR41/43 are more prevalent in hypertensive patients. Our findings highlight that lack of SCFA-receptor signalling via both GPR41/43 increases risk of high blood pressure, suggesting these receptors could be targeted as a new treatment.

## Main text

High blood pressure is the leading risk factor for death, accounting for one in every five deaths worldwide.^1^ Uncontrolled high blood pressure leads to stroke, coronary heart disease, chronic kidney disease, and dementia.^2^ Since the 1970s, researchers have shown that a diet high in fibre lowers blood pressure,^3, 4^ but the underlying mechanisms remain unclear. Understanding these mechanisms allows for the development of new therapies.

Fermentable dietary fibre feeds the commensal microbiota in the large intestine, producing the short-chain fatty acids (SCFAs) acetate, propionate and butyrate as by- products.^5^ Although these metabolites are found in the host’s systemic circulation, these metabolites are almost exclusively produced by the gut microbiota.^6^ In both the intestine and systemically, SCFAs are potent regulators of the host’s metabolism and immune function.^7^ We and others previously showed that a high fibre diet or supplementation with SCFAs reduced blood pressure and its accompanying cardio-renal complications in experimental models of hypertension^8–11^ and, most recently, in human hypertension.^12^ In addition, SCFAs influence epithelial cell proliferation, differentiation, and repair, thereby modulating gut barrier integrity.^13^ Impaired gut barrier integrity allows the translocation of detrimental substances, such as lipopolysaccharides (LPS), from the gastrointestinal tract into the peripheral circulation,^14^ activating a cascade of pro-inflammatory events. Chronic pro-inflammatory processes have been demonstrated to increase blood pressure and are a hallmark of cardiovascular disease.^15^

SCFAs activate cell signalling in the host via G-protein coupled receptors. The two most abundant receptors that sense SCFAs, GPR41 and GPR43, are expressed in immune cells, such as macrophages, and cells in the gastrointestinal tract (Extended Data Figure 1). They can be activated by all three main SCFAs (Figure 1a) and have redundant roles in several pathways.^16^ Animals individually lacking GPR41 or GPR43 have some form of cardiovascular dysfunction, but their effect on blood pressure is unclear.^8, 17^ Importantly, hypertensive patients have lower expression of *GPR43* mRNA in peripheral blood cells.^18^ However, whether the blood pressure-regulating effect of SCFAs depends on these receptors is unknown.

**Figure 1.**
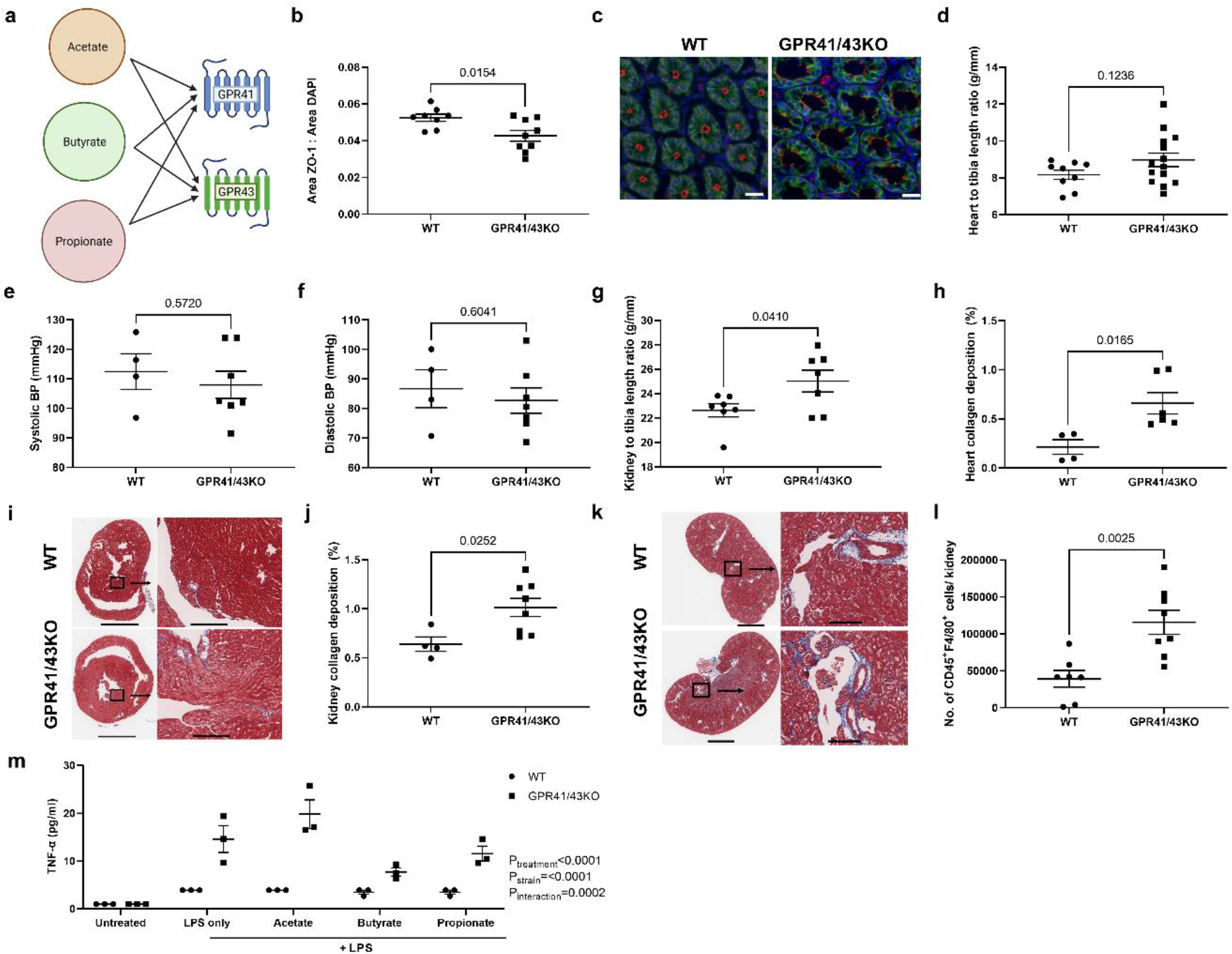
Baseline characteristics of GPR41/43KO and WT mice. a, Brief schematic diagram showing acetate, butyrate, and propionate binding to both GPR41 and GPR43. **b,** Zonulin-1 (ZO-1) expression as a ratio of area of ZO-1 staining/area of 4′,6-diamidino-2- phenylindole (DAPI) staining; and **c,** representative images of composite ZO-1, DAPI, epithelial cell adhesion molecule (EPCAM) staining in the colon (scale bars 20µm, 40X magnification). **d,** Heart weight to tibia length ratio in g/mm; **e,** Systolic and **f,** Diastolic blood pressure in mmHg; **g,** Kidney weight to tibia length ratio in g/mm; **h,** Heart collagen deposition as a percentage of the region of interest and **I,** representative micrographs of Masson’s trichrome staining indicative of fibrosis in heart (scale bars 2mm and 200µm respectively); **j,** Kidney collagen deposition as a percentage of the region of interest and **k,** Representative micrographs of Masson’s trichrome staining indicative of fibrosis in the kidney (scale bars 3mm and 200µm respectively); l, Number of CD45^+^F4/80^+^ cells per kidney; **m**, Levels of the pro-inflammatory cytokine TNF-α produced by in vitro stimulation of bone marrow derived macrophages of GPR41/43KO and WT mice treated with lipopolysaccharides (LPS) only or in combination with acetate, butyrate, or propionate. Panels b,d,e-h,j,l; unpaired two-tailed t-test. Panel m; 2-way ANOVA with p-values adjusted for FDR. Data shown as mean +/- SEM, n=3-14.

Considering the redundancy in pathways activated by GPR41 and GPR43, we hypothesised that mice lacking both GPR41 and GPR43 would have a more severe hypertensive phenotype when challenged to develop hypertension and would remain unprotected against cardiovascular disease when fed a high-fibre diet. Here we aimed to mechanistically determine whether the cardiovascular protective effect of SCFAs in hypertension is dependent on these receptors. We employed a unique whole-body double knockout (KO) lacking GPR41/43 to assess their role in blood pressure and cardiorenal regulation. Our results in mice indicate that a high-fibre diet can attenuate hypertension by improving gut barrier integrity and reducing systemic inflammation via macrophages, and that this is dependent on the SCFA receptors GPR41 and GPR43. We then used a translational approach using the UK Biobank to determine that the genetic predisposition to lower expression of both GPR41 and GPR43 increases the risk of hypertension.

## Results

### Gut barrier integrity is disrupted in GPR41/43KO mice

SCFAs reduced expression of protein kinase C (PKC) proteins in human intestinal epithelial cells *in vitro.*^19^ In rat intestinal cells, one of PKC family members, PKCα, increased phosphorylation and expression of zonulin-1 (ZO-1), a key epithelial tight junction protein that maintains the gut epithelial barrier.^20^ This process may be dependent on GPR43 and GPR41.^21^ Thus, we first hypothesised that naïve GPR41/43KO mice would exhibit impaired gut barrier integrity. Using immunofluorescence, we found that, relative to wild-type (WT) mice, GPR41/43KO mice had significantly lower ZO-1 protein expression in the colon (Figure 1b-c, Extended Data Figure 2).

**Figure 2.**
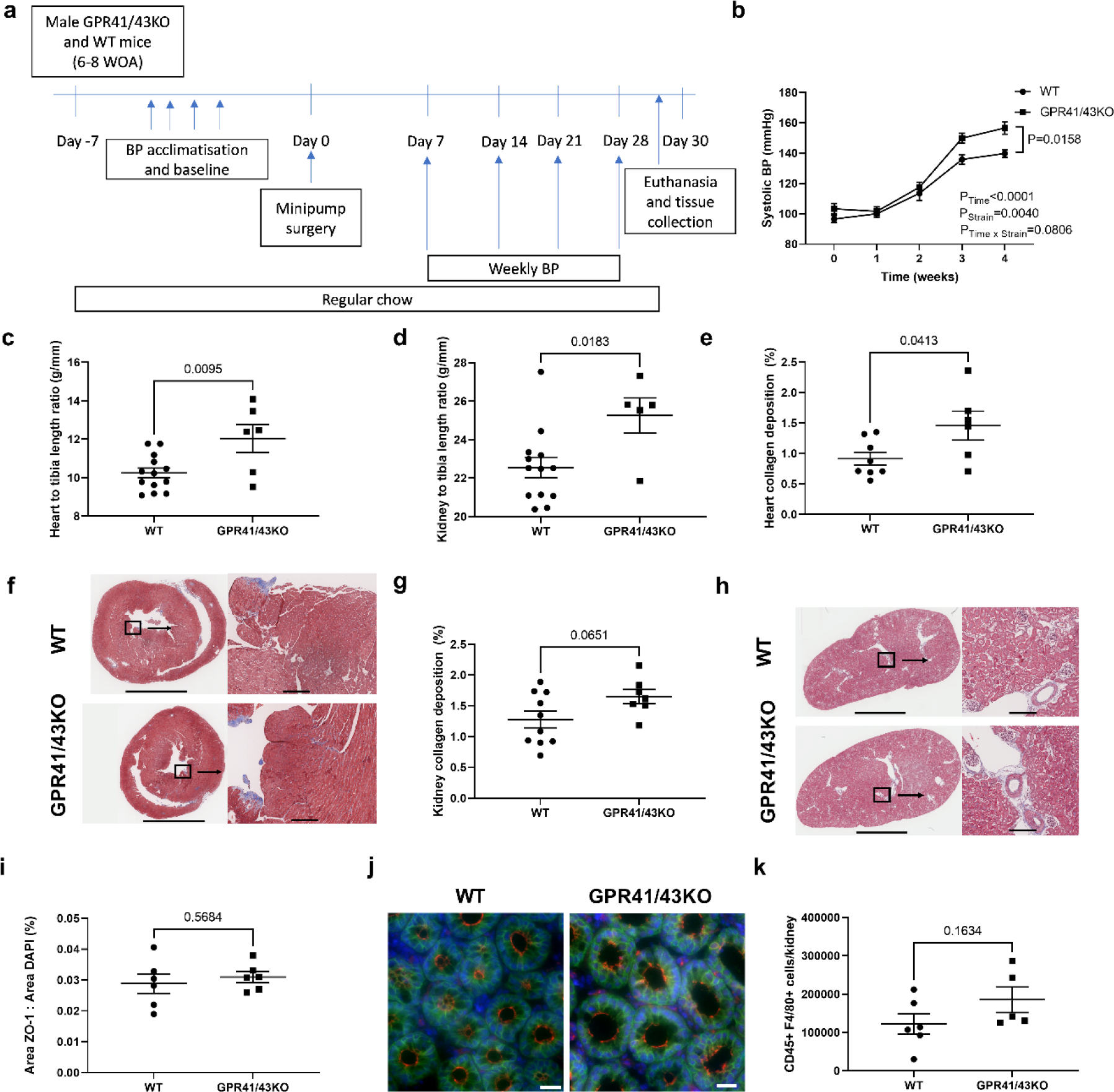
Angiotensin II treatment of GPR41/43KO mice and WT mice. a, Brief schematic of the study design. Male GPR41/43 knockout (KO) and wildtype (WT) mice at 6- 8 weeks of age underwent blood pressure (BP) acclimatisation and baseline measurement.Mice remained on regular chow through the study period. Mice underwent minipump surgery containing vehicle or Angiotensin II under isoflurane anaesthesia. Following minipump surgery, weekly BP recording was performed. Mice were euthanised by CO2 asphyxiation at the end of the 4-week protocol. Tissues were collected for further analysis; **b**, Systolic BP in mmHg; **c,** Heart and **d,** Kidney weight to tibia length ratio in g/mm; **e,** Heart collagen deposition as a percentage of the region of interest; **f**, Representative micrographs of Masson’s trichrome staining indicative of fibrosis in the heart (scale bars 2mm and 200µm respectively); **g**, Kidney collagen deposition as a percentage of the region of interest; **h**, Representative micrographs of Masson’s trichrome staining indicative of fibrosis in the kidney (scale bars 3mm and 200µm respectively; **i**, Zonulin-1 (ZO-1) expression as a ratio of area of ZO-1 staining/area of 4′,6-diamidino-2-phenylindole (DAPI), staining in the colon; **j**, Representative images of composite ZO-1, DAPI, epithelial cell adhesion molecule (EPCAM) staining in the colon (scale bars 20µm, 40X magnification); **k**, Number of CD45^+^F4/80^+^ cells per kidney; Panel b; 2-way ANOVA with p-values adjusted for FDR. Panels c-e, g, i, k; unpaired two-tailed t-test. Data shown as mean +/- SEM, n=5-13.

### Naïve GPR41/43KO mice have more cardio-renal fibrosis

Naïve adult GPR41/43KO mice had a similar heart weight to tibia length ratio (Figure 1d) and blood pressure (Figure 1e-f), but they had significantly larger kidneys (Figure 1g) and higher collagen deposition levels in the heart (Figure 1h-i) and kidney (Figure 1j-k) compared to WT mice. We next hypothesised that increased gut permeability would allow gut microbiota-derived toxins, such as LPS, to enter the host’s peripheral circulation, activating immune cells such as macrophages, leading to inflammation. Indeed, we found significantly higher F4/80^+^ macrophages in the kidney of GPR41/43KO mice compared to WT mice (Figure 1l and Extended Data Figure 3), suggesting the activation of pro-inflammatory pathways, which are present in hypertension.^22^

**Figure 3.**
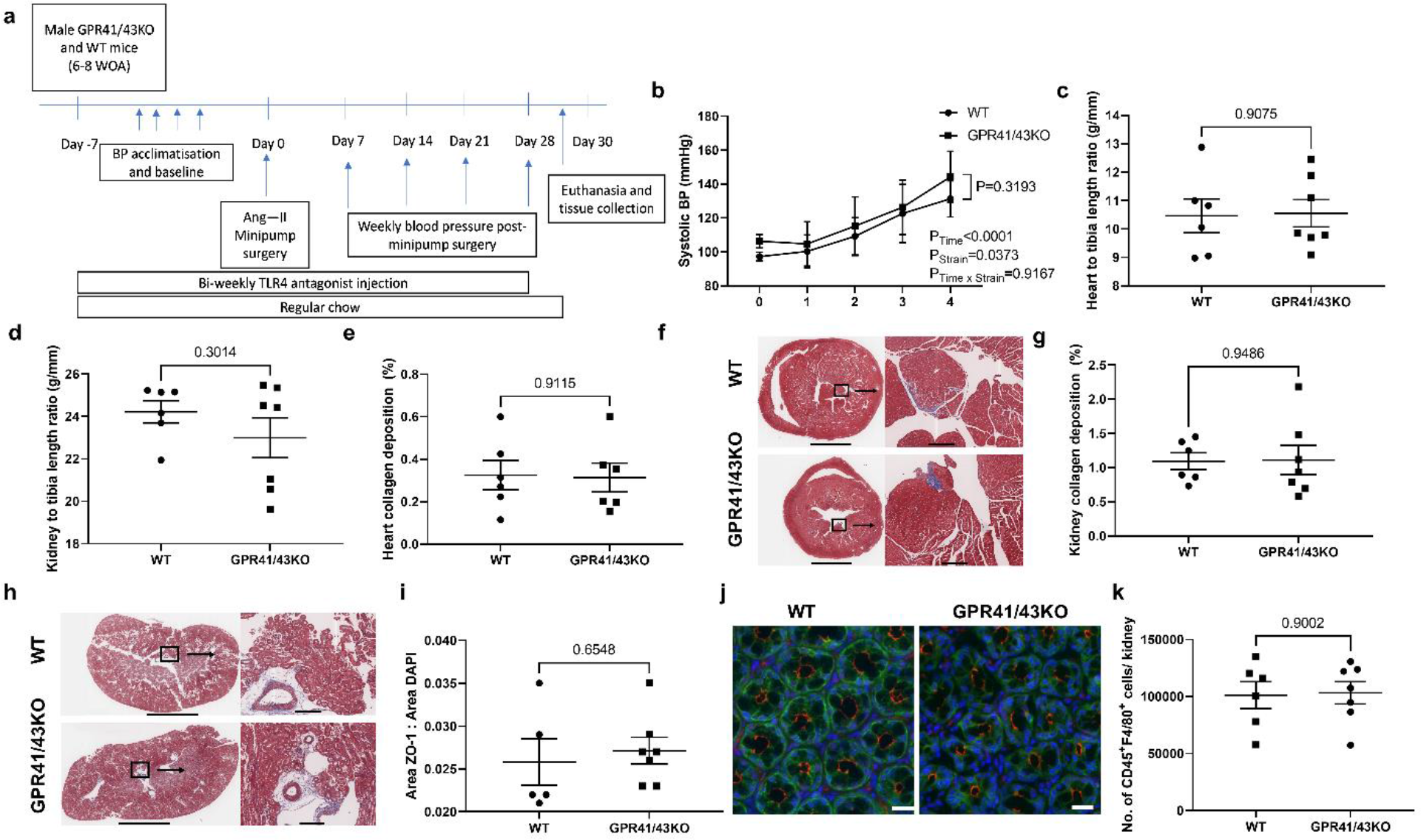
TLR4 antagonist treatment rescues phenotype of angiotensin II GPR41/43KO mice and WT mice. a, Brief schematic of the study design. Male GPR41/43 knockout (KO) and wildtype (WT) mice at 6-8 weeks of age underwent blood pressure (BP) acclimatisation and baseline measurement. Mice remained on regular chow through the study period and received biweekly intraperitoneal injections of TLR4 antagonist, lipopolysaccharide from *Rhodobacter sphaeroides* (LPS-RS), from one week before surgery until the endpoint euthanasia and tissue collection. Mice underwent minipump surgery containing Angiotensin II (Ang II) under isoflurane anaesthesia. Following minipump surgery, weekly BP recording was performed. Mice were euthanised by CO2 asphyxiation at the end of the 4-week protocol. Tissues were collected for further analysis; **b,** Systolic BP in mmHg ; **c,** Heart and **d,** Kidney weight to tibia length ratio in g/mm; **e,** Heart collagen deposition as a percentage of the region of interest ; **f,** Representative micrographs of Masson’s trichrome staining indicative of fibrosis in the heart (scale bars 2mm and 200µm respectively) ; **g,** Kidney collagen deposition as a percentage of the region of interest; **h,** Representative micrographs of Masson’s trichrome staining indicative of fibrosis in the kidney (scale bars 3mm and 200µm respectively); **i,** Zonulin-1 (ZO-1) expression as a ratio of area of ZO-1 staining/area of 4′,6- diamidino-2-phenylindole (DAPI) staining in the colon; **j,** Representative images of composite ZO-1, DAPI, epithelial cell adhesion molecule (EPCAM) staining in the colon (scale bars 20µm, 40X magnification); **k,** Number of CD45^+^F4/80^+^ cells per kidney. Panel b, 2-way ANOVA with p-values adjusted for FDR. Panels c-e, g, i, k; unpaired two-tailed t-test. Data shown as mean +/- SEM, n=5-7.

In addition to regulating gut barrier integrity, SCFA-receptors are highly expressed by immune cells such as macrophages (Extended Data Figure 1b).^23^ Studies show macrophages in the kidney produce inflammatory cytokines leading to the development of tissue fibrosis,^24, 25^ which could explain the worsened fibrosis observed in the GPR41/43KO mice.

To confirm whether macrophages from GPR41/43KO mice are more sensitive to LPS, we stimulated bone marrow-derived macrophages (BMDMs) from naïve GPR41/43KO and WT mice with SCFAs and LPS. We found that BMDMs from GPR41/43KO mice were non-responsive to SCFAs and produced significantly higher pro-inflammatory cytokine TNF-α levels (Figure 1m).

### Hypertensive GPR41/43KO mice have a worse cardiovascular phenotype

As a result of the baseline phenotype, we hypothesised that, compared to WT mice, GPR41/43KO mice would have a more severe phenotype when challenged with a hypertensive stimulus, such as Angiotensin II (Ang II, Figure 2a). We found that GPR41/43KO mice treated with Ang II for 4-weeks had significantly higher blood pressure, heart and kidney weights, and kidney and heart fibrosis levels (Figure 2b-h). There was no difference in the ZO-1 expression in the colon and F4/80^+^ kidney macrophages of Ang II treated GPR41/43KO and WT mice (Figure 2i-k). However, Ang II-treatment resulted in lower ZO-1 expression in the colon of both GPR41/43KO and WT mice, irrespective of genotype (Extended Data Figure 4a), confirming previous findings that hypertension leads to impaired gut barrier integrity.^26^ Hypertensive GPR41/43KO and WT mice also had significantly higher F4/80^+^ macrophages in the kidney (Extended Data Figure 4b), suggesting that Ang II leads to a phenotype similar to lack of signalling via GPR41/43.

**Figure 4.**
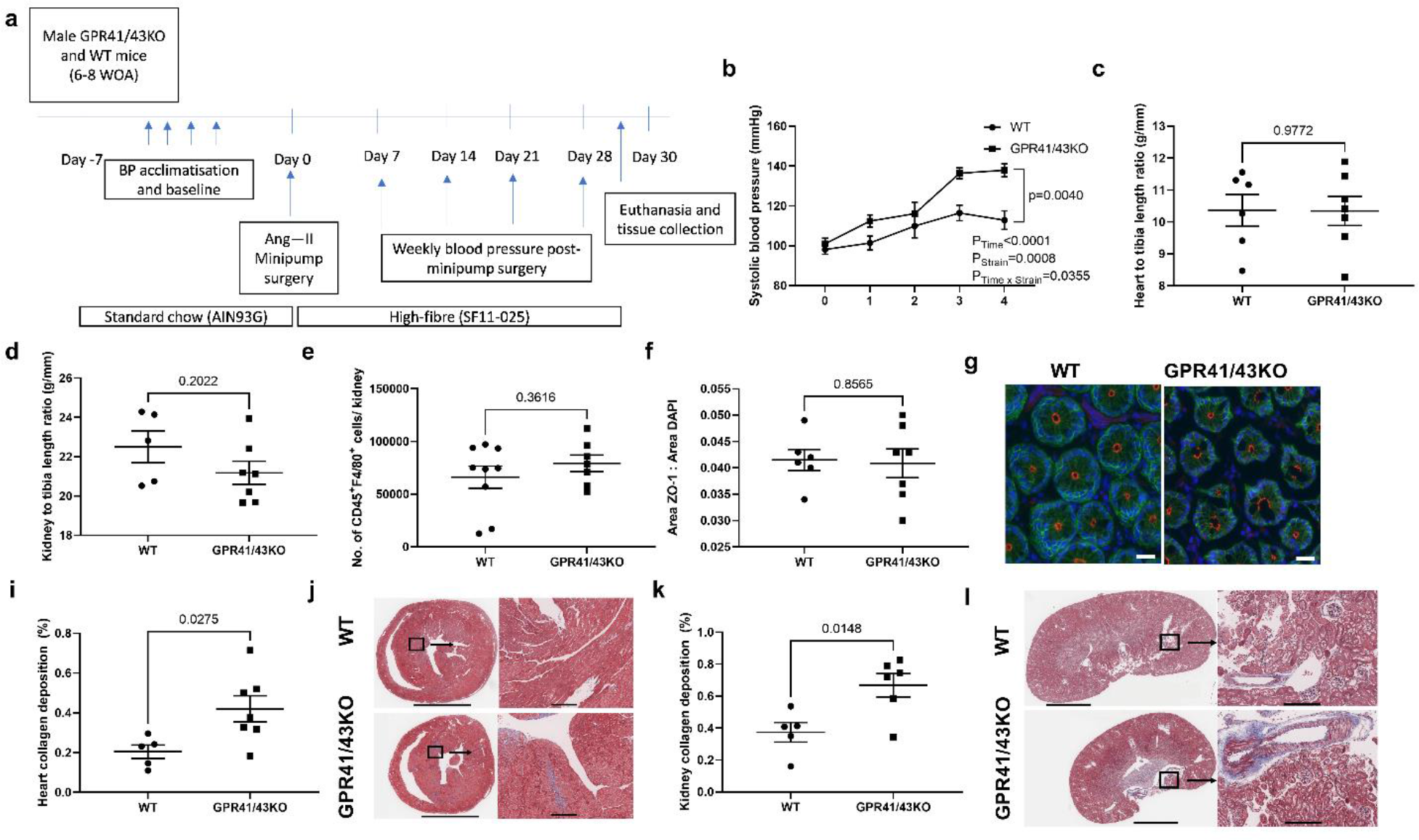
A high-fibre diet rescues phenotype of Angiotensin II treated GPR41/43KO mice and WT mice. a, A brief schematic of the study design. Male GPR41/43 knockout (KO) and wildtype (WT) mice at 6-8 weeks of age underwent blood pressure (BP) acclimatisation and baseline measurement. Mice were fed a control diet (AIN93G) for 1 week prior to minipump surgery. Mice underwent minipump surgery containing Angiotensin II (Ang II) under isoflurane anaesthesia. Following minipump surgery, mice were switched to a high-fibre diet, SF11-025 (on the AIN93G background). Weekly BP recording was performed. Mice were euthanised by CO2 asphyxiation at the end of the 4-week protocol. Tissues were collected for further analysis; **b,** Systolic BP in mmHg; **c,** Heart and **d,** Kidney weight to tibia length ratio in g/mm; **e,** Number of CD45^+^F4/80^+^ cells per kidney; **f,** Zonulin- 1 (ZO-1) expression as a ratio of area of ZO-1 staining/area of 4′,6-diamidino-2-phenylindole (DAPI) staining in the colon; **g**, Representative images of composite ZO-1, DAPI, epithelial cell adhesion molecule (EPCAM) staining in the colon (scale bars 20µm, 40× magnification); **h,** Heart collagen deposition as a percentage of the region of interest; **i,** Representative micrographs of Masson’s trichrome staining indicative of fibrosis in the heart (scale bars 2mm and 200µm respectively); **j,** Heart collagen deposition as a percentage of the region of interest; **k,** Representative micrographs of Masson’s trichrome staining indicative of fibrosis in the kidney (scale bars 3mm and 200µm respectively). Panel b, 2-way ANOVA with p-values adjusted for FDR. Panels c-f, h,j; unpaired two-tailed t-test. Data shown as mean +/- SEM, n=5-7.

### Blocking TLR4 improves the cardiovascular phenotype in GPR41/43KO mice

The reduced gut barrier integrity and the presence of higher number of macrophages in a distal organ such as the kidney led us to hypothesise that GPR41/43KO mice would have detectable levels of LPS in the peripheral circulation. We first performed an ELISA to quantify LPS in the plasma of Ang II-treated GPR41/43KO and WT mice, but levels were below the detection threshold (data not shown). Considering LPS ELISA assays may not be appropriate outside the setting of septicaemia,^27^ we quantified activation of the LPS receptor TLR4 in the plasma using the HEK-Blue TLR4 assay. We discovered that Ang II treatment increased TLR4 activation in both GPR41/43KO and WT mice (Extended Data Figure 5).

**Figure 5.**
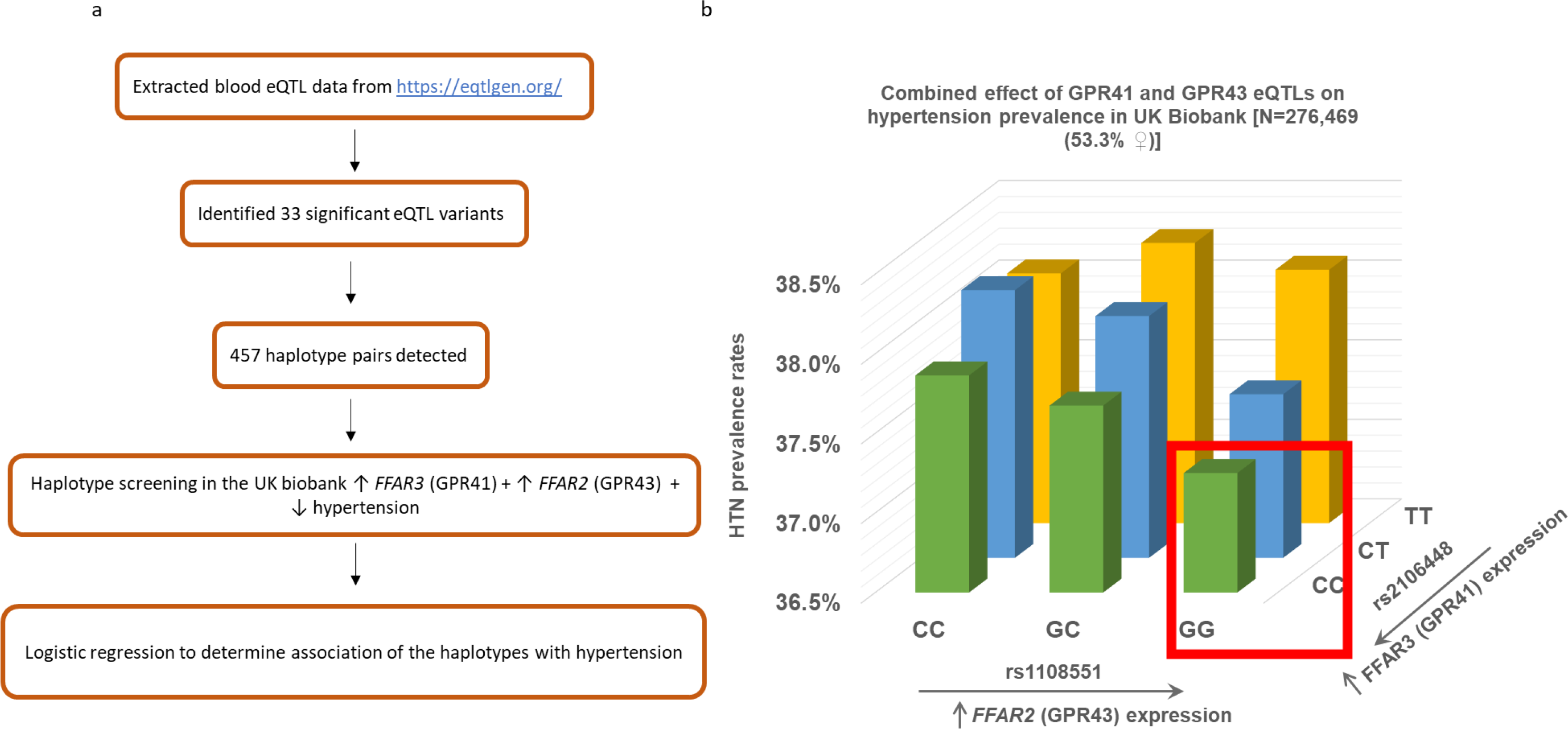
**Combined increased expression of both GPR41 and GPR43 is protective against hypertension. a**, Brief schematic for the identification and association of combined *FFAR3* (GPR41)/*FFAR2* (GPR43) expression in hypertensive individuals. Expression quantitative loci (eQTL) of GPR41 and GPR43 were extracted from eQTLgen. A total of 33 significant eQTL variants with 457 haplotype pairs detected. Haplotype screening was then performed in the UK Biobank which fit with the desired phenotype - increase in GPR41 and GPR43 expression and lower hypertension prevalence. Logistic regression was then performed to determine the association of the haplotypes with hypertension. **b,** Hypertension prevalence rates of participants in the UK Biobank (n=276,469) with combined expression quantitative loci (eQTL) in rs1108551 near *FFAR2* (GPR43) and rs2106448 near *FFAR3* (GPR41) genes. eQTL variants were estimated and tested for its association with hypertension using a logistic regression model with Plink v1.9, including age, sex, and BMI as covariates.

We then treated hypertensive GPR41/43KO and WT mice with a TLR4 antagonist to determine if we could rescue the hypertensive phenotype (Figure 3a). TLR4 antagonist treatment abolished the cardiovascular differences previously observed between WT and GPR41/43KO mice, which had similar blood pressure, kidney and heart weight to tibia length ratio , and fibrosis (Figure 3b-h). Moreover, there was no difference in the ZO-1 expression in the colon and F4/80^+^ macrophages in the kidney between GPR41/43KO and WT mice after TLR4 antagonist treatment (Figure 3i-k). This shows that blocking TLR4 signalling restored ZO-1 expression and reduced activation of systemic macrophages, providing indirect evidence that LPS spill over from the gut microbiome into the systemic circulation might be responsible for the cardiovascular phenotype observed when there is a lack of signalling via GPR41/43.

### A high fibre diet protects against cardiovascular disease via GPR41/43 signalling

We next hypothesised that a high-fibre diet would attenuate hypertension by improving gut barrier function, thereby preventing LPS translocation from the gut into the systemic circulation. To determine if the protective effects of a high fibre diet depend on GPR41/43 signalling, we fed a high fibre diet to GPR41/43KO and WT mice and challenged them with Ang II (Figure 4a). The high-fibre diet maintained blood pressure around baseline in WT mice, but not in the GPR41/43KO mice (Figure 4b), but the increase in blood pressure was lower than in control diet-fed KO mice (Extended Data Figure 6a). There was no difference between high fibre-fed GPR41/43KO and WT mice in terms of organ weights, renal F4/80^+^ macrophages and ZO-1 expression levels in the colon, but GPR41/43KO mice still had higher fibrosis levels in the heart and kidney relative to WT mice (Figure 4c-l). In addition, we compared regular chow-fed and high fibre-fed mice, and observed a significant increase in ZO-1 expression in the colon independent to strain (Extended Data Figure 2,6b), confirming our hypothesis that overall high fibre diet improves gut barrier function. High fibre diet also significantly reduced the numbers of F4/80^+^ macrophages in hypertensive kidneys, independent of strain (Extended Data Figure 6c). We also compared the colonic ZO-1 levels between high fibre-fed mice and TLR4 antagonist-treated mice, and found that ZO-1 levels were significantly higher in mice fed high fibre, suggesting that high fibre is protective by improving gut barrier integrity and not merely blocking LPS translocation (Extended Data Figure 2, 7).

### SNPs in eQTL for the genes GPR41/GPR43 are associated with human hypertension

Finally, we aimed to determine if GPR41/43 expression is associated with human hypertension. To determine the combined effects of GPR41/43, we first extracted the expression quantitative trait loci (eQTL) for both GPR41 (*FFAR3* gene) and GPR43 (*FFAR2* gene) in peripheral blood mononuclear cells available from a public database.^28^ We identified 277 significant eQTLs (FDR<0.05) for the *FFAR3* (GPR41) and *FFAR2* (GPR43) genes. We identified 33 significant eQTL variants following linkage disequilibrium (LD) pruning, and then performed haplotype estimation and association in the UK Biobank cohort (n=276,469 after quality control, Extended Data Table 1). Haplotypes were tested at two variant levels for a combined effect that would lower both GPR41 and GPR43. In total, 457 haplotypes were detected, and we performed association tests in the same regression model like the individual variant. Since we hypothesised that the combined increased expression of both GPR41 and GPR43 would be protective against hypertension, haplotypes were screened to identify those with the same association direction (i.e., if increased eQTL of both genes was associated with protection against hypertension). We selected the pair with the lowest *P*-value to validate in the UK Biobank. We found that participants with the G allele in the SNP rs2106448 near the *FFAR3* (upstream intergenic variant) and the C allele in the SNP rs1108551 near the *FFAR2* (downstream intergenic variant) gene, associated with higher expression of these genes, had a significantly lower prevalence of hypertension (37.3% in homozygous carriers of both alleles compared to 38% in the rest of the cohort, *P*=0.00763) (Figure 5b). This is the first translational evidence that suggests that GPR41/43 signalling might be relevant in human hypertension.

## Discussion

Our study provided mechanistic insights into how a high fibre diet confers protection in hypertension and cardiovascular disease. The absence of the receptors GPR41 and GPR43 in hypertensive mice lead to significantly higher tissue inflammation, blood pressure, and kidney and heart collagen deposition levels. Using a TLR4 antagonist, which improved gut barrier integrity, we likely reduced the translocation of LPS from the gut into the host’s systemic circulation, thereby inhibiting TLR4 activation and subsequent inflammation-induced cardiorenal damage and fibrosis. We also demonstrated that GPR41/43 partially mediated the protective effects of a high fibre diet in hypertension but did not mediate the improvement in gut barrier integrity. Finally, in a large human cohort, we provide evidence that the combined effect of two eQTLs that increase the expression of the genes coding for GPR41 and GPR43 is associated with lower prevalence of hypertension.

Targeting the gut and its microbiome for therapeutic approaches, such as for cardiovascular disease prevention and treatment, has been challenging due to the lack of understanding of the causative aspects of the gut microbiota and/or their interactions with the host.^29^ Here we leveraged our understanding of the host response to SCFA signalling and sought to manipulate the host’s physiology instead. Previous studies showed pathophysiological changes in the gut, including lower tight junction protein expression in spontaneously hypertensive rats and hypertensive patients^26, 30^ and higher plasma LPS levels in hypertensive patients.^26^ Our findings are the first functional evidence suggesting that improving the gut barrier integrity is a potential therapeutic option for lowering blood pressure. We further demonstrate that gut barrier integrity in a hypertensive state depends on a high-fibre diet and SCFAs.

An unanswered question in our study is the importance of SCFA-receptor signalling in immune compared to non-immune compartments in hypertension. Our findings at present point out that both compartments play a role. Previous studies showed that GPR41 and GPR43 signalling in both epithelial and immune cells were responsible for the phenotype observed in colitis.^31, 32^ Our findings suggest that lack of GPR41 and GPR43 leads to impaired gut barrier integrity. Bone-marrow chimaeras and specific knockout of GPR41 and GPR43 in gut epithelial and immune cells would further deepen our understanding of how SCFAs regulate systemic physiological responses in the host.

Another major issue in the microbiome field has been the lack of translation of findings from experimental models to humans. For example, the use of SCFAs to lower blood pressure in treatment-naïve hypertensive patients was only recently demonstrated.^12^ Consistently with a multifactorial contribution to blood pressure regulation, here we provide translational evidence that the combined higher expression of GPR41/43 may have a small effect in decreasing the prevalence of hypertension. Genome-wide association studies performed in the past have identified variants in these genes that were associated with hypertension; however, they did not reach genome-wide statistical significance.^33^ The common database used to extract eQTL data is the Genotype-Tissue Expression (GTEx); however, we did not discover any significant eQTLs for GPR41/43 genes in the kidney, heart, and whole blood (data not shown). Therefore, we extended our search to other databases available. The eQTLGen Consortium comprises 37 available datasets with a total of 31,684 individuals, much larger than the GTEx database, with samples from 1,000 individuals. However, the eQTLGen Consortium has data from blood samples only. We know that immune cells in the blood express GPR41/43 (Extended Data Figure 1), so we decided to determine whether there was an association between the combined expression of GPR41/43 in blood cells and hypertension. Although we found that the combined expression of GPR41/43 in the blood cells was associated with lower hypertension prevalence, eQTL data from other organs, including the heart, kidneys, and blood vessels such as the aorta, would further expand our understanding of the tissue-specific roles of these receptors in cardiovascular health.

We expected the Ang II treated GPR41/43KO to have lower ZO-1 expression and higher kidney macrophage infiltration than WT. However, this was not the case. This suggests that Ang II treatment copies a phenotype driven by lack of GPR41/43, as these were observed in naïve KO mice. We also observed only a partial rescue in the high-fibre diet fed hypertensive GPR41/43KO mice further suggesting other mechanisms are at play. We speculate that other SCFA receptors, such as GPR109A (a non-classical SCFA receptor), may also be involved in this improved gut epithelial barrier.^34^ In addition, we have recently demonstrated that SCFA production lowers colonic pH and activates GPR65, which also had partial cardiovascular protection, likely via immune-driven mechanisms.^35^ Thus, our findings highlight the extensive redundancy involved in SCFA signalling, which is not surprising given how important fibre and SCFAs are for several health aspects.^36^ Other KO models of SCFA receptors and specific agonists targeting SCFA receptors would further expand our understanding and confirm whether they are potential therapeutic targets. Finally, while there are sex differences in response to Ang II,^37^ we acknowledge as a limitation that we only studied male mice. However, the human cohort (n=276,469) included 53.3% female participants, demonstrating that our findings may still be relevant to blood pressure regulation in females.

In conclusion, we show that signalling via the SCFA-receptors GPR41/43 attenuate hypertension by improving gut barrier integrity and preventing the translocation of LPS and activation of TLR4. This mechanism opens the possibility of targeting these receptors as a novel treatment for hypertension and other diseases where gut barrier integrity is impaired.

## Methods

### Animal experiments

We obtained animal ethics approval from the Monash University Animal Ethics Committee (AEC 17465 and AEC 27929). Whole-body single-knockout GPR41 and GPR43 mice colonies were established by C.R.M. at Monash University. GPR41/43 double KO mice colony was generated by using CRISPR/Cas9-based protocol at the Monash Genome Modification Platform (MGMP) at Monash University. Briefly, the UCSC Genome Browser was used to identify guide RNA target sites flanking the Gpr41/43 genes. The following guide RNA were used: 868 bp upstream of the ATG of Gpr41 (5’ TACCTGTAACCCAGACGTTA 3’) and 1145 bp downstream of the STOP codon of Gpr43 (5’ AAGCCAGCTACGGGCTACAC 3’) to knock out both Gpr41 and Gpr43 genes.

CRISPR RNAs (crRNA, IDT) were annealed with trans-activating crRNA (tracrRNA) to form a functional crRNA:tracrRNA guide RNA duplex. Cas9 nuclease (IDT, # 1081058) was incubated with the guide RNAs to form a ribonucleoprotein (RNP) complex. Cas9 nuclease (30ng/μl) and the crRNA:tracrRNA guide RNA duplexes (30ng/μl) were microinjected into the pronucleus/cytoplasm of the zygotes at the pronuclei stage. Injected zygotes were transferred into the uterus of pseudo pregnant F1 females. Genome-edited F1 GPR41/43 KO were bred with C57BL/6 mice for two cycles to dilute the off-target effects. Homozygous littermates were generated by breeding heterozygous parents. These mice were maintained as a homozygous breeding colony and genotyped periodically. Age-matched WT mice C57BL/6J MARP were obtained from the Monash Animal Research Platform and bred in the same facility under similar conditions to the GPR41/43KO colony.

### Dietary interventions

Diets used in this study include i) regular chow (Barastoc rat and mouse pellets; Crude fibre ∼6%; Ridley); ii) standard chow (AIN93G, Crude fibre ∼4.7%; Speciality Feeds); iii) diet rich in resistant starches (high resistant starch, HF, SF11-025; Crude fibre ∼9.7%; Speciality Feeds). The diet rich in resistant starches is referred to as high-fibre diet.

### Blood pressure measurement

Systolic and diastolic blood pressure were measured using a non-invasive tail-cuff machine (CODA, Kent Scientific Corporation). Mice were acclimatised for three days before baseline blood pressure measurement. For mice that underwent vehicle or Ang II surgery, weekly blood pressure was recorded for four consecutive weeks starting one week after surgery.

Following 10 minutes of acclimatisation to the restrainer, blood pressure recordings were performed for 15 cycles (the first five cycles were omitted as part of acclimatisation). At least four recordings were averaged to calculate systolic and diastolic blood pressure.

### Osmotic minipump surgery (subcutaneous)

Six to eight-week-old male WT and GPR41/43KO mice were randomised using Excel random sort function and underwent subcutaneous osmotic minipump (Alzet 2004) insertion surgery under isoflurane anaesthesia. Mice received minipump containing either vehicle (0.9% saline) or Ang II (Auspep) at 0.5 mg/kg body weight/day for four weeks.

### TLR4 antagonist treatment

Mice received intraperitoneal injections of ultrapure LPS from *Rhodobacter sphaeroides* (LPS-RS, Invivogen), a potent TLR4 antagonist (50µg/kg body weight dose as per^38^). Mice received biweekly injections starting from one week before minipump surgery until the week of endpoint tissue collection (10 injections in total).

### Endpoint tissue collection

At endpoint, mice were euthanised by CO2 asphyxiation. Following exsanguination and perfusion with Dulbecco’s phosphate buffered saline (dPBS, Thermo Scientific), organs were collected and sectioned if necessary. Samples for tissue fibrosis analyses were stored in 10% neutral buffered formalin for 24-48 hours before paraffin-embedding step. Samples for immunofluorescence studies were stored in OCT embedded cassettes, and some tissue in dPBS on ice for flow cytometry studies which requires fresh tissue samples.

### Cardiac and renal collagen quantification

Mouse heart and kidney were fixed in 10% neutral buffered formalin for 24-48 hours before being paraffin-embedded and sectioned at 4μm by the Monash Histology Platform. Masson’s trichrome staining was then performed to quantify collagen deposition in the organs. Total collagen in the heart and kidney sections was quantified using a colour-threshold macro developed by Dr Chad Johnson using ImageJ. The operator was blinded of the strain and treatment groups. Collagen levels were expressed as a percentage of the region of interest.

Scale bars are shown in each figure and explained in respective figure legends.

### Isolation of mouse immune cells from kidney and spleen

One kidney from each mouse was mechanically dissociated using scissors and digested with 0.156mg/ml Collagenase type XI (Sigma-Aldrich), 0.030mg/ml Hyaluronidase type IV-S (Sigma-Aldrich), and 1.8mg/ml Collagenase type I (Sigma-Aldrich). Following digestion, samples were filtered and underwent 40/80% Percoll (Sigma-Aldrich) separation. The mononuclear cells at the interface were collected, washed, and underwent FACS staining.

Mouse spleen cells were isolated for use as single colour controls for flow cytometry. Mouse spleen tissue were filtered, and red blood cell lysed.

### Flow cytometry

An entire kidney and 10^6^ spleen cells were used for FACS staining. Kidney and spleen cells were first stained for viability using LIVE/DEAD Fixable Violet Dead Cell Stain kit (Life Technologies) and stained with anti-CD45 and anti-F4/80 (rat anti-mouse, Biolegend) and then fixed and permeabilised using the FOXp3/Transcription Factor Staining Buffer Set (eBioscience). Kidney cell counts were determined using flow cytometry counting beads (CountBright Absolute, Life Technologies) and adjusted per kidney. Fixed and stained samples were acquired within 24-48 hours. Samples were acquired using a five-laser BD LSRFortessa X-20 flow cytometer (Monash FlowCore Facility). Analysis was then performed using FlowJo (Tree Star). We adhered to the guidelines and best practices of using flow cytometry set here.^39^

### Immunofluorescence

We performed immunofluorescence staining for ZO-1 as previously described.^40^ Colon sections were obtained during endpoint tissue harvest, splayed open, and embedded in OCT- cassettes using OCT and dry ice and then stored at -80°C until staining. In brief, we stained 10-micron colon sections for ZO-1 (Invitrogen, 1:100), EpCAM (Invitrogen, 1:500), and DAPI (Invitrogen). ZO-1 expression was quantified using representative images. Six images per section were taken blindly. Image J was used to quantify the area of ZO-1 relative to the area of DAPI staining.

### Isolation, culture, and stimulation of bone-marrow-derived macrophages

As previously described, we performed isolation and culture of bone-marrow-derived macrophages (BMDMs).^41^ In brief, isolated bone marrow cells from the tibia and femur of WT and GPR41/43KO mice at 8-12 weeks of age and were flushed using a 25G needle. 10^7^ bone marrow cells were plated in a tissue culture treated plate. Cells were cultured inL929 supplemented media (DMEM high glucose with 10% FBS) to initiate differentiation. After overnight incubation in a tissue-culture-treated plate, non-adherent cells were transferred to a non-tissue-culture treated plate and cultured for four days. BMDMs were then lifted using Trypsin (Sigma-Aldrich), and plated at a density of 50,000 cells per well in a 24-well plate in 500µl of media without L929 supplementation. We added 1mM SCFAs (Sigma Aldrich) acetate, propionate, and butyratefor 30 minutes, before adding LPS from Escherichia coli (Sigma Alrich) at 10ng/ml for 20 hours.

### HEK-Blue TLR4 assay

HEK-Blue TLR4 assay was performed as previously described^42^. In brief, HEK-Blue TLR4 cells or HEK-Blue null cells (Greiner, 1 x 10^5^ in 200 µl culture media) were seeded in 96- well plates until 80-90% confluence. The cells were then either stimulated with 0.9% normal saline (negative control), 6.25ng/ml LPS (positive control), or 50µl plasma samples from the treatment groups (Sham and Ang II treated WT and GPR41/43KO mice). After 24 hours, 20µl of supernatant were transferred to a fresh 96-well plate and incubated with 180µl of QUANTI-Blue solution (Invivogen) at 37°C. SEAP activity was measured at 625 nm using a CLARIOstar plate reader (BMG Labtech).

### Enzyme-linked immunosorbent assay (ELISA)

Concentrations of tumour necrosis factor (TNF)-α from tissue culture supernatants post-stimulation were measured using ELISA MAX Standard Set Mouse TNF-α kit (Biolegend) according to the manufacturer’s protocol.

### Statistical analyses of animal experiments

We used GraphPad Prism (version 9) for all statistical analyses. We first tested and determined that the data were normally distributed using Shapiro-Wilk test. When two groups were compared, independent unpaired two-tail t-test was used. Two-way analysis of variance (2-way ANOVA) with two-stage step-up method of Benjamini, Krieger and Yekutieli’s false discovery rate adjustment for multiple comparisons was used to compare between the various treatment groups and strains. For systolic blood pressure data, we used mixed effects analysis with two-stage step-up method of Benjamini, Krieger and Yekutieli’s false discovery rate adjustment for multiple comparisons. All values are presented as mean +/- SEM, and two-tail P<0.05 were considered significant.

#### Blinding statement

Operators were blinded to treatment groups during acquisition of data for in vivo (tail-cuff measurement) and ex vivo experiments (immunofluorescence, ELISAs, TLR4 assay, fibrosis quantification, and flow cytometry).

### Expression quantitative trait loci (eQTL) in GPR41 and GPR43 genes

Expression quantitative trait loci (eQTL) in GPR41 and GPR43 genes were extracted from eQTLGen Consortium^28^, which collected associations between genotypes and whole-blood-derived expressions from 31,684 individuals. After filtering for FDR<0.05 and pairwise linkage disequilibrium (pruning at r2=0.2 by Plink v1.9^43^), 33 significant independent eQTL associations (28 in GPR41 and 5 in GPR43) were selected for following analyses.

### Human study cohort

UK Biobank (UKBB) is a large general population cohort with available genotype data and health-related information from approximately half a million individuals. This research has been conducted using the UK Biobank Resource under Application Number 86879. We applied a stringent quality control pipeline to remove samples with genetic non-Caucasians, genotype-phenotype sex discrepancies, or kinship to other participants, resulting in a total of 276,469 individuals. Individuals’ hypertension phenotype was defined according to their electronic medical records and the10^th^ Revision of the International Classification of Diseases (ICD10) code ‘I10’, or their self-reported previous doctor’s diagnosis of ‘Essential hypertension’ from the verbal interview. Each haplotype from individual GPR41 and GPR43 eQTL variants were estimated and tested for its association with hypertension using a logistic regression model with Plink v1.9^43^, including age, sex, and BMI as covariates. Only haplotypes with P-value < 0.05 were kept for further investigation. Statistical computing environment R (v3.6.0) was used for processing UK Biobank data. Genotypes visualization of selected eQTL variants was produced by Microsoft Excel (v16.66.1)

## Data availability

The data underlying this article can be shared for selected research questions upon reasonable request to the corresponding author – please email F.Z.M. at, who will respond within 4 weeks.

## Code availability

Plink v1.9: https://www.cog-genomics.org/plink/ and R: https://www.r-project.org/. The full code can be shared upon reasonable request to the corresponding author – please email F.Z.M. at, who will respond within 4 weeks.

## Acknowledgements

We would like to acknowledge the Monash Histology Facility for support with histology, the Monash FlowCore facility for support with flow cytometry, Monash Animal Research Platform for support with animal work, the Monash Bioinformatics Platform for access to M3 servers, and Dr Dominic de Nardo for the L929 cells. This research has been conducted using the UK Biobank Resource under Application Number 86879. This work was supported by a National Health & Medical Research Council (NHMRC) of Australia Project Grant (GNT1159721), and fellowship to D.M.K. F.Z.M. is supported by a Senior Medical Research Fellowship from the Sylvia and Charles Viertel Charitable Foundation, a National Heart Foundation Future Leader Fellowship (105663), and NHMRC Emerging Leader Fellowship (GNT2017382). R.R.M. is supported by a scholarship from the Faculty of Science, Monash University. L.X. is supported by a Monash Graduate Scholarship. J.O.D is supported by NHMRC ECR Fellowship (GNT 1124288). The Baker Heart & Diabetes Institute is supported in part by the Victorian Government’s Operational Infrastructure Support Program.

## Author Contribution Statement

R.R.M. planned and performed most of the in vivo and in vitro experiments, data analyses, provided intellectual inputs and wrote the manuscript. T.Z. performed analysis involving eQTLs and validation in the U.K Biobank cohort, E.D. provided major contribution to animal in vivo and in vitro experiments; A.B.W., L.X., H.J., E.S., M.P. and M.N. assisted with assisted with in vivo animal experiments; C.J. developed the macro used for the fibrosis quantification; N.B. and M.K.L. performed the TLR4 cell-based assay; D.M.K. provided intellectual input and secured funding; J.O.A. assisted with flow-cytometry experiments, provided intellectual input, and supervised the study; C.R.M. developed the knockout mice colonies, provided intellectual input, and supervised the study; F.Z.M. designed the study, secured funding, provided intellectual input, and supervised the study. All authors had full access to all data in the study, revised the manuscript critically, approved the version to be published, and had the final responsibility for the decision to submit the manuscript for publication.

