## Supplementary figures and tables for "GPR41/43 regulates blood pressure by improving gut epithelial barrier integrity to prevent TLR4 activation and renal inflammation"

[1]Hypertension Research Laboratory, School of Biological Sciences, Monash University, Melbourne, Australia; [2]Institute for Medical Research, Ministry of Health Malaysia, Kuala Lumpur, Malaysia; [3]Bioimaging Platform, La Trobe University, Melbourne Australia; [4]Department of Microbiology, Monash Biomedicine Discovery Institute, Monash University, Melbourne, Australia; [5]Monash Bioimaging Facility, Monash University, Melbourne, Australia; [6]Department of Microbiology, Anatomy, Physiology and Pharmacology, La Trobe University, Melbourne, Australia; [7]La Trobe Centre for Extracellular Vesicles, La Trobe University, Melbourne, Australia; [8]Heart Failure Research Group, Baker Heart and Diabetes Institute, Melbourne, Australia; [9]Department of Cardiology, Alfred Hospital, Melbourne, Australia; [10]Central Clinical School, Faculty of Medicine Nursing and Health Sciences, Monash University, Melbourne, Australia; [11]School of Pharmaceutical Sciences, Shandong Analysis and Test Center, Qilu University of Technology (Shandong Academy of Sciences), Jinan, 250014, China.

***Corresponding author**: A/Prof Francine Marques, Hypertension Research Laboratory, School of Biological Sciences, Faculty of Science, Monash University, Melbourne, Australia, Phone: +61-03-9905 6958.

#### Extended Data Table

**Extended Data Table 1. Demographics and clinical characteristics of participants.**

| Variable | Normotensives | Hypertensives | *P*-value* |
|---|---|---|---|
| **Sample size (n, %)** | 171,488 | 104,981 | |
| **Age (years)** | 55.0 (±8.0) | 59.8 (±7.0) | <0.0001 |
| **BMI (kg/m$^2$)** | 26.3 (±4.2) | 29.0 (±5.1) | <0.0001 |
| **Sex (% female)** | 57.6 | 46.3 | <0.0001 |
| **SBP (mmHg)** | 124.5 (±39.8) | 139.8 (±42.5) | <0.0001 |
| **DBP (mmHg)** | 73.7 (±24.3) | 80.3 (±25.2) | <0.0001 |
| **BP-lowering medication (%)** | 0.7 | 30.4 | <0.0001 |

Data are shown as mean ± standard deviation or numbers and percentages. Legend: body mass index, BMI; systolic blood pressure, SBP; diastolic blood pressure; waist to hip ratio, WHR. *P-values are calculated from Students' t tests for continuous variables or Chi-squared tests for categorical variables

### Extended Data Figures

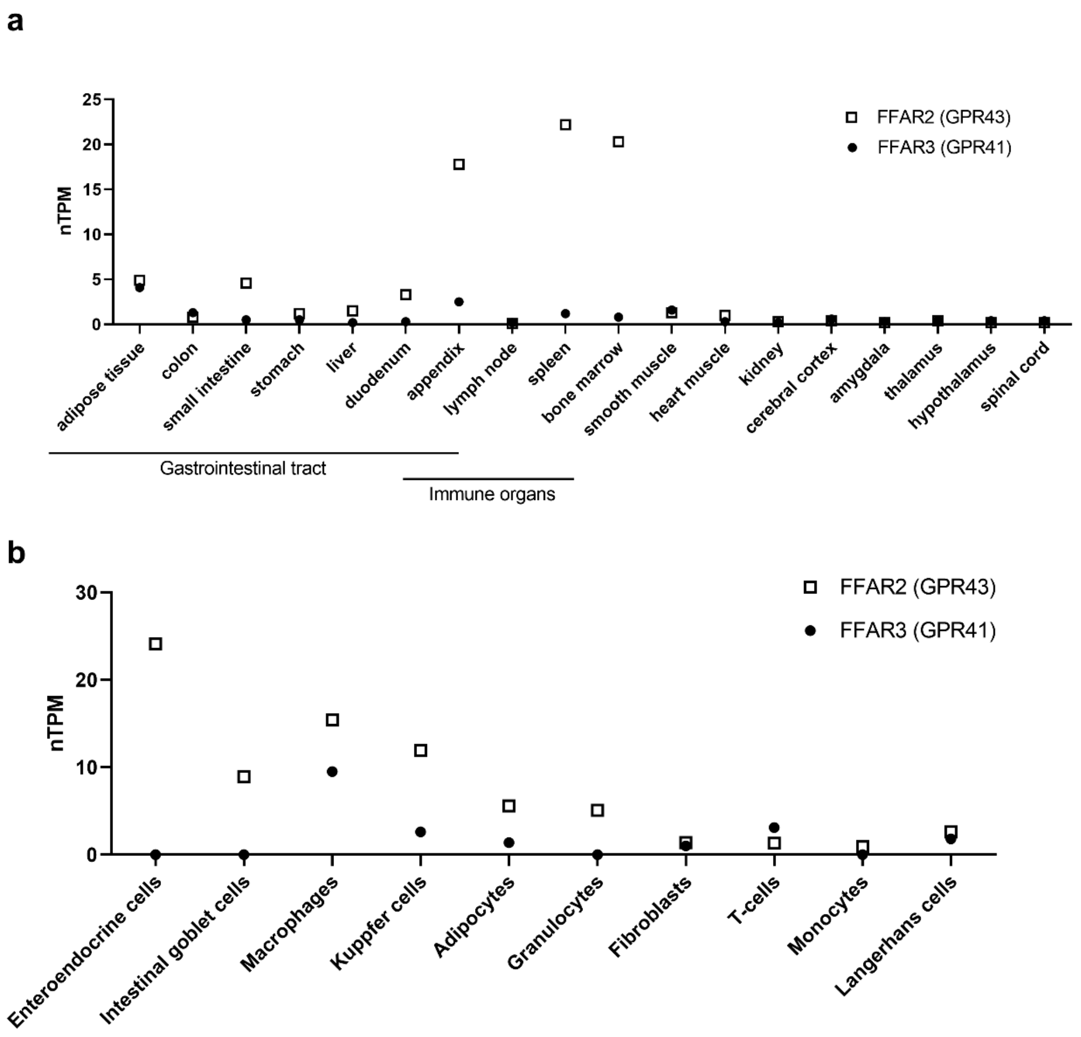


**Extended Data Figure 1.** GPR41 (*FFAR3*) and GPR43 (*FFAR2*) expression in **a,** tissue and **b,** single-cell, expressed as normalized transcripts per million (nTPM). Data from the Human Protein Atlas (www.proteinatlas.org).

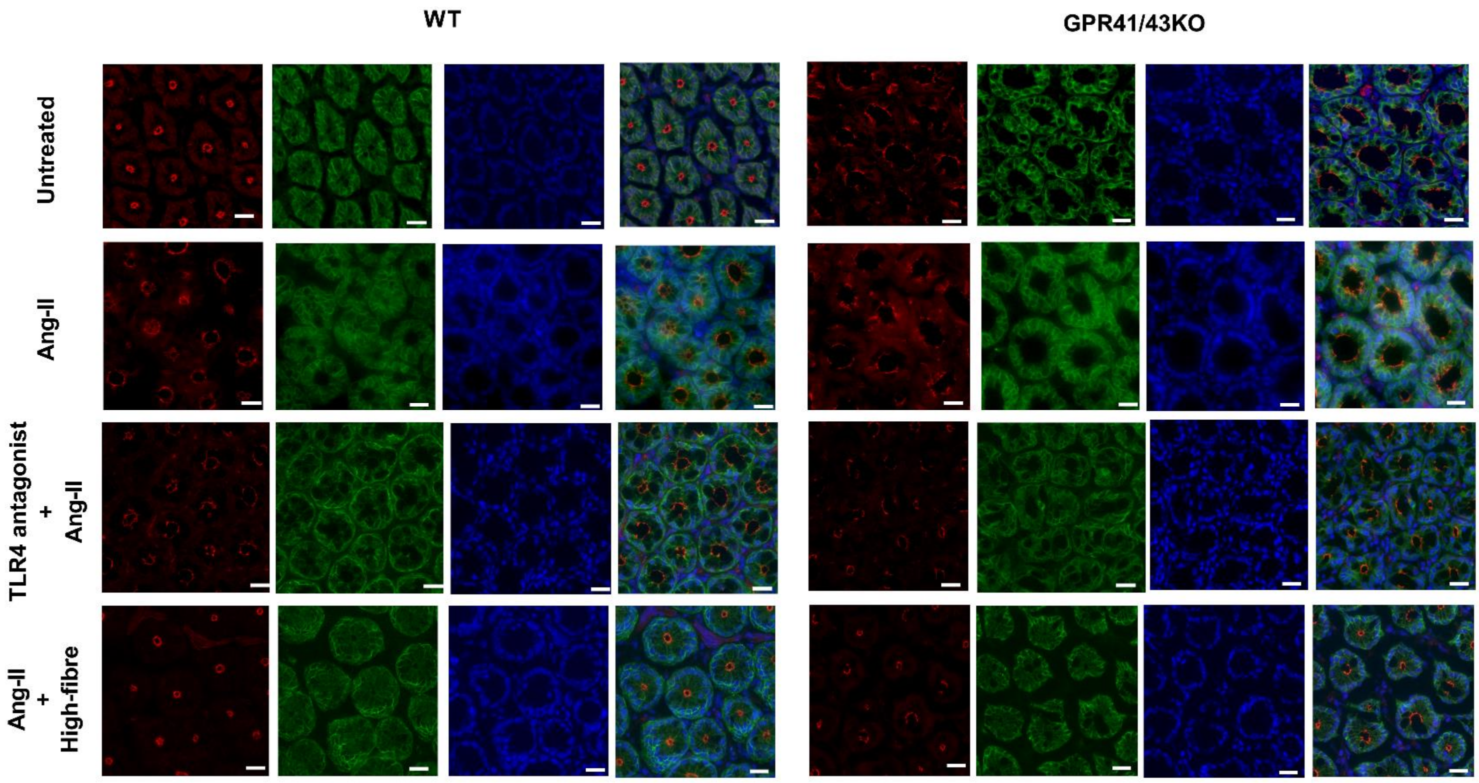


**Extended Data Figure 2.** Representative images of individual zonulin-1 (ZO-1), epithelial cell adhesion molecule (EPCAM), 4′,6-diamidino-2-phenylindole (DAPI) and merged composite channels of GPR41/43KO and wildtype (WT) mice of various treatment groups. (scale bars 20µm, 40X magnification). Angiotensin II (Ang II).

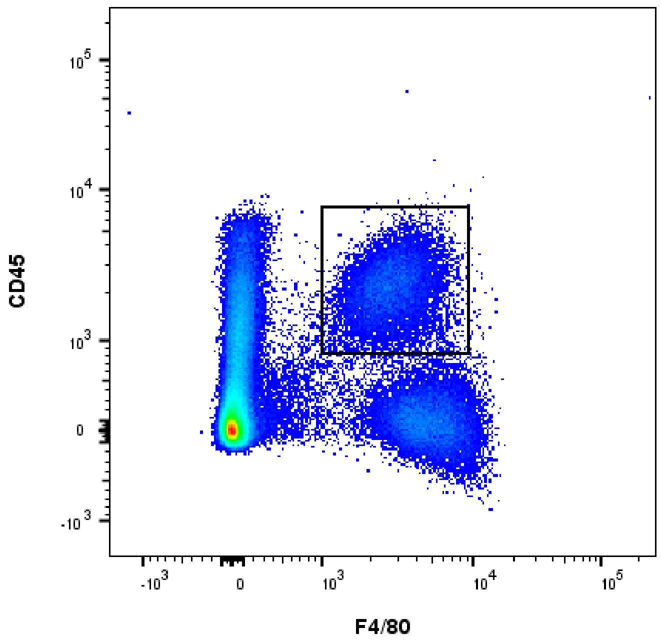


**Extended Data Figure 3**. Gating strategy for flow cytometry of CD45$^+$F4/80$^+$ cells kidney macrophages.

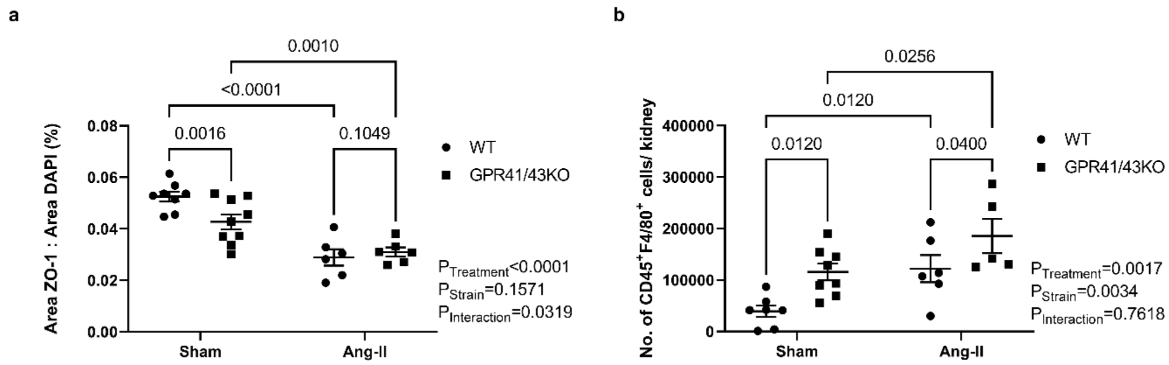


**Extended Data Figure 4. Angiotensin II treatment leads to lower expression of Zonulin-1 and increases kidney macrophage numbers. a,** Zonulin-1 (ZO-1) expression as a ratio of area of ZO-1 staining/area of 4′,6-diamidino-2-phenylindole (DAPI) staining, and **b,** the number of $CD45^{+}F4/80^{+}$ cells kidney macrophages in sham and Angiotensin II (Ang II) treated GPR41/43 knockout (KO) and wildtype (WT) mice, 2-way ANOVA with p-values adjusted for FDR, data shown as mean +/- SEM, n=6-9.

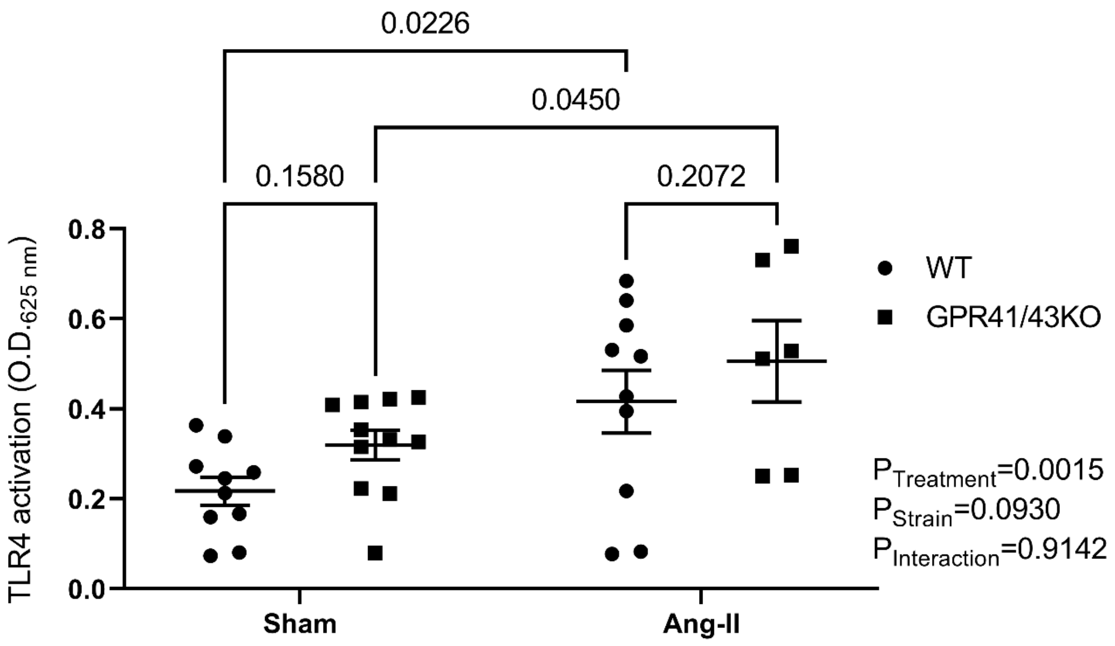


**Extended Data Figure 5. TLR4-activation levels in plasma of sham and Angiotensin II (Ang II) treated GPR41/43 knockout (KO) and wildtype (WT) mice**, 2-way ANOVA with *P*-values adjusted for FDR, data shown as mean +/- SEM, n=6-11.

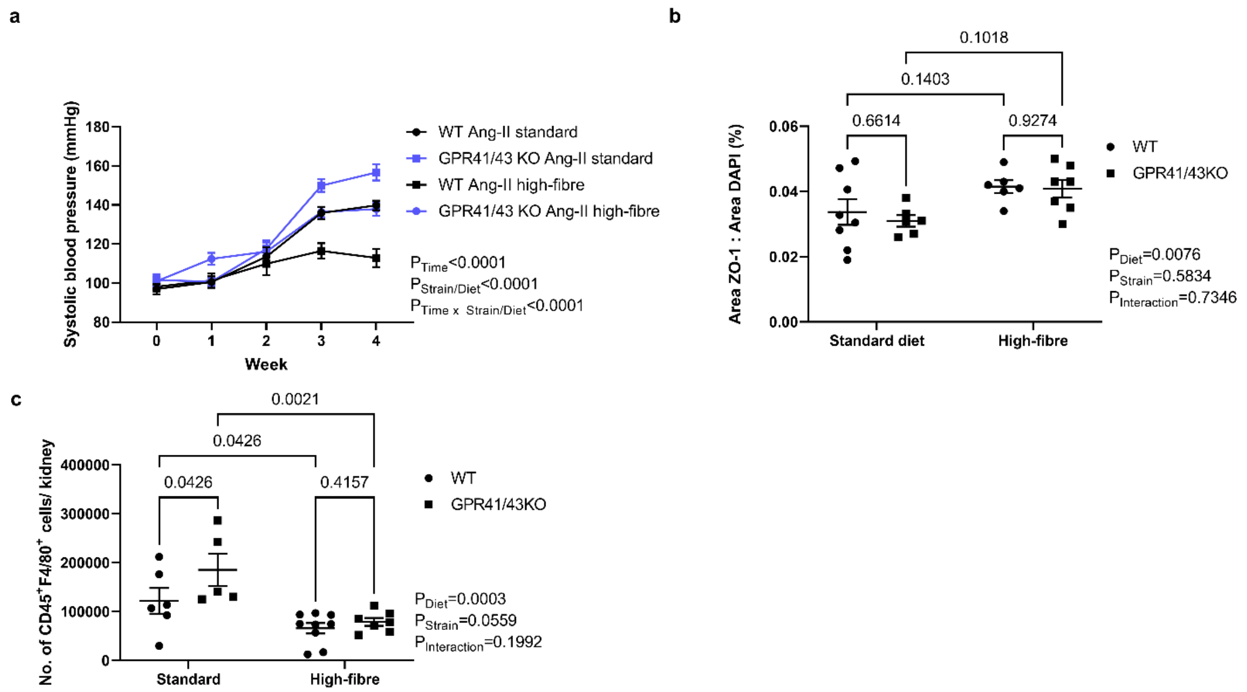


**Extended Data Figure 6**. **A high-fibre diet increases Zonulin-1 expression, decreases macrophage numbers in the kidney, and influences systolic blood pressure**. **a,** systolic blood pressure (BP) of standard chow and high-fibre fed Angiotensin II (Ang II) treated GPR41/43 knockout (KO) and wildtype (WT) mice; **b,** Zonulin-1 (ZO-1) expression as a ratio of area of ZO-1 staining/area of 4′,6-diamidino-2-phenylindole (DAPI) staining, and **c,** the number of $CD45^{+}F4/80^{+}$ cells per kidney , 2-way ANOVA with p-values adjusted for FDR, data shown as mean +/- SEM, n=5-9.

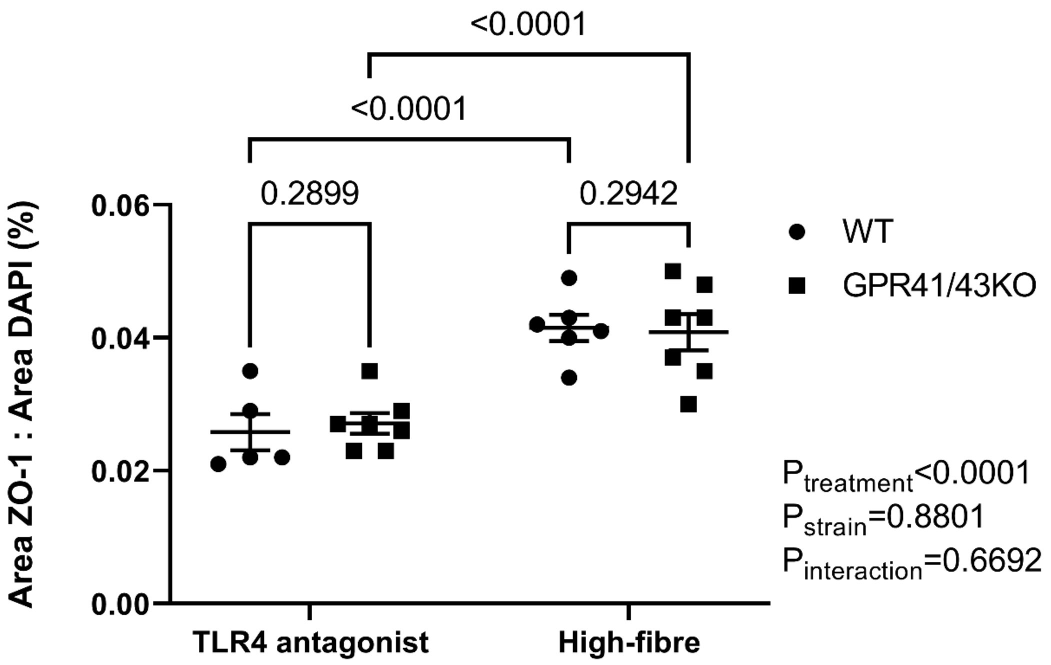


**Extended Data Figure 7.** Zonulin-1 (ZO-1) expression as a ratio of area of ZO-1 staining/area of 4′,6-diamidino-2-phenylindole (DAPI) staining of hypertensive TLR4 antagonist treated and high-fibre fed GPR41/43 knockout (KO) and wildtype (WT) mice, 2-way ANOVA with p-values adjusted for FDR, data shown as mean +/- SEM, n=5-7.
